# Neutralizing Antibodies Isolated by a site-directed Screening have Potent Protection on SARS-CoV-2 Infection

**DOI:** 10.1101/2020.05.03.074914

**Authors:** Xiaoyu Liu, Fang Gao, Liming Gou, Yin Chen, Yayun Gu, Lei Ao, Hongbing Shen, Zhibin Hu, Xiling Guo, Wei Gao

## Abstract

Neutralizing antibody is one of the most effective interventions for acute pathogenic infection. Currently, over three million people have been identified for SARS-CoV-2 infection but SARS-CoV-2-specific vaccines and neutralizing antibodies are still lacking. SARS-CoV-2 infects host cells by interacting with angiotensin converting enzyme-2 (ACE2) via the S1 receptor-binding domain (RBD) of its surface spike glycoprotein. Therefore, blocking the interaction of SARS-CoV-2-RBD and ACE2 by antibody would cause a directly neutralizing effect against virus. In the current study, we selected the ACE2 interface of SARS-CoV-2-RBD as the targeting epitope for neutralizing antibody screening. We performed site-directed screening by phage display and finally obtained one IgG antibody (4A3) and several domain antibodies. Among them, 4A3 and three domain antibodies (4A12, 4D5, and 4A10) were identified to act as neutralizing antibodies due to their capabilities to block the interaction between SARS-CoV-2-RBD and ACE2-positive cells. The domain antibody 4A12 was predicted to have the best accessibility to all three ACE2-interfaces on the spike homotrimer. Pseudovirus and authentic SARS-CoV-2 neutralization assays showed that all four antibodies could potently protect host cells from virus infection. Overall, we isolated multiple formats of SARS-CoV-2-neutralizing antibodies via site-directed antibody screening, which could be promising candidate drugs for the prevention and treatment of COVID-19.

## Introduction

Coronavirus disease 2019 (COVID-19) is a worldwide epidemic of respiratory disease caused by the novel human coronavirus SARS-CoV-2^1,2^. Currently, over three million infected people have been identified in more than 200 countries and regions by laboratory testing, with an average mortality rate of approximately 6% (https://covid19.who.int/). The real number of infected cases is even higher, considering the detection limitation in many counties. Therefore, there is an urgent need to develop an effective vaccine and neutralizing antibody against SARS-CoV-2.

SARS-CoV-2, a single-stranded positive-sense RNA virus of the β-*Coronaviridae* family^3^. It shares 79% nucleotide sequence identity with SARS-CoV-1^4^. Both SARS-CoV-2 and SARS-CoV-1 infect host cells by directly interacting with the host angiotensin-converting enzyme-2 (ACE2) receptor through their spike glycoprotein expressed on the viral membrane and subsequently trigger the fusion of the cell and virus membrane for cell entry^5,6^. Spike glycoprotein exists as a homotrimeric complex on the viral membrane of coronaviruses^7^. Each spike monomer contains an S1 subunit and an S2 subunit^8^. The S1 subunit binds to ACE2 through its receptor-binding domain (RBD) to initiate cell recognition, whereas the S2 subunit anchors the spike protein to the viral envelope and responds to S1-induced cell recognition to mediate effective membrane fusion via a conformational transition^7^. These determined infection mechanisms indicated that blocking the interaction of SARS-CoV-2-RBD and ACE2 would cause a direct neutralizing effect against virus.

Neutralizing antibody is one of the most effective interventions for acute pathogenic infection^9^. Several approaches are reported to obtained SARS-CoV-2 neutralizing antibodies successfully. Among them, one approach is to screen the preexisting SARS-CoV-1 antibody repertoires by evaluating cross-reactivity^10^. An alternative approach is to clone the neutralizing antibody from the isolated SARS-CoV-2-RBD-specific single B cells from infected patients^11,12^. However, the feasibility of these two strategies is quite limited due to the rare chance of accessing either SARS-CoV-1 antibodies or SARS-CoV-2-infected patients. Therefore, *in vitro* site-directed screening in a human antibody library would be more feasible and efficient.

The RBD is a relatively isolated domain of the S1 subunit with independent function^13^. The crystal structure of the SARS-CoV-2-RBD^14^ and the SARS-CoV-2-RBD/ACE2 complex^15,16^ has already been reported. It presents quite similar interaction details compared to those of the previously determined SARS-CoV-1-RBD/ACE2 structure^17^. Notably, a recent study performed a systematic bioinformatics analysis to predict the potential B cell epitope and T cell epitope of SARS-CoV-2^18^. The only predicted conformational B cell epitope in the RBD is located within the ACE2 interface (P491-Y505). This information suggests that the ACE2 interface of SARS-CoV-2-RBD might have high immunogenicity, which would be a suitable targeting epitope to develop SARS-CoV-2-specific antibodies with potent neutralizing function by *in vitro* screening.

In the current study, we selected the ACE2-interface of SARS-CoV-2-RBD as the targeting epitope to screen neutralizing antibody. We performed site-directed antibody screening by phage display and finally obtained one IgG antibody and three single domain antibodies with potent neutralizing activities for SARS-CoV-2. These neutralizing antibodies are promising candidate drugs for the prevention and treatment of COVID-19.

## Materials and Methods

### Plasmids and reagents

The coding sequences of ACE2 and the SARS-CoV-2 spike were cloned into the pLVX vector to construct stable cell lines. The coding sequences of truncated ACE2 (Q18-S740), SARS-CoV-1-RBD (P317-V510), SARS-CoV-2-RBD (P330-V524) and SARS-CoV-2-RBD mut were cloned into the pFUSE vector to obtain hFc-fusion proteins. SARS-CoV-2-RBD mut was designed by substituting key residues on SARS-CoV-2-RBD with Ala or Phe to disrupt its interaction with ACE2 (Table 1). The coding sequences of domain antibodies were also cloned into the pFUSE vector to obtain hFc-fusion proteins. The coding sequences of the 4A3 heavy chain variable region and light chain variable region were amplified by adding the IL-2 signal peptide and cloned into the expression vectors pFUSE-CHIg-HG1 and pFUSE2-CLIg-hk (Invivogen, San Diego, CA), respectively. All plasmids were identified by sequencing. SARS-CoV-2-RBD-his protein was purchased from GenScript (GenScript, Nanjing), and GPC5-his protein was purchased from R&D (Minneapolis, MN).

**Table 1.** Mutated residues of SARS-CoV-2-RBD mut.

| SARS-CoV-1 | SARS-CoV-2 | SARS-CoV-2 mut |
| --- | --- | --- |
| K 390 | R 403 | A 403 |
| D 392 | D 405 | A 405 |
| V 404 | K 417 | A 417 |
| Y 436 | Y 449 | A 449 |
| Y 440 | Y 453 | A 453 |
| P 470 | E 484 | A 484 |
| L 472 | F 486 | A 486 |
| N 473 | N 487 | A 487 |
| Y 475 | Y 489 | A 489 |
| N 479 | Q 493 | A 493 |
| D 480 | S 494 | A 494 |
| G 482 | G 496 | F 496 |
| Y 484 | Q 498 | A 498 |
| T 486 | T 500 | A 500 |
| T 487 | N 501 | A 501 |
| Y 491 | Y 505 | A 505 |

### Structural modeling

SARS-CoV-2-RBD/ACE2 complex (PDB ID: 6M0J), ectodomain of SARS-CoV-2 spike timer (closed state) (PDB ID: 6VXX) and ectodomain of SARS-CoV-2 spike timer (open state) (PDB ID: 6VYB) were downloaded from the RCSB Protein Data Bank. The structure model of SARS-CoV-2-RBD mut was predicted by the Protein Fold Recognition Server Phyre2. The remodeled ectodomain trimers of SARS-CoV-2 spike (open state) and SARS-CoV-2 spike (closed state) were established by replacing the partially determined RBD of the ectodomain trimers of SARS-CoV-2 spike (open state) (PDB ID: 6VYB) and SARS-CoV-2 spike (closed state) (PDB ID: 6VXX) with the completely determined SARS-CoV-2-RBD (PDB ID: 6M0J) by using PyMOL, Discovery Studio and SWISS MODEL.

### Phage display

The TG1 clone was picked and cultured overnight in 2YT medium at 37 °C. SARS-CoV-2-RBD-his or SARS-CoV-2-RBD-hFc proteins in PBS buffer were coated on ELISA plates at 4 °C overnight. The phage library (Tomlison I library and Domain antibody library) and the coated wells were blocked with PBST with 5% milk at room temperature for 1 h. The blocked phages were precleaned by negative antigens and then added into the coated wells to incubate for 1 h at room temperature. After washing 20 times with PBST, the SARS-CoV-2-RBD-binding phages were eluted with 100 mM triethylamine (Sigma-Aldrich, Xuhui, Shanghai). All eluted phages were collected and used to infect TG1 cells. After incubating with helper phages, the eluted phages were rescued with a titer of approximately 10^11^~10^12^ pfu/ml for the next round of screening.

### ELISA

For direct ELISA, the indicated antigen (5 μg/ml) was coated on an ELISA plate at 4 °C overnight. After blocking, biotin-labeled ACE2, blocked phage or the indicated antibodies were added to the wells and incubated at 37 ° C for 0.5 h.

Streptavidin-HRP (Thermo, Pudong New Area, Shanghai), a rabbit anti-M13 HRP antibody (for phage) (GE Healthcare, Milwaukee, WI), or a goat anti-human Fcγ HRP antibody (Jackson ImmunoResearch, West Grove, PA) was added. TMB and H_2_SO_4_ were added to detect the OD_450_ nm value. For capture ELISA, an anti-his antibody (5 μg/ml) was coated on an ELISA plate at 4 °C overnight. After blocking, soluble antibodies extracted from the periplasm of TG1 were added into the wells and incubated at 37 °C for 0.5 h. After washing, SARS-CoV-1-RBD-hFc, SARS-CoV-2-RBD-hFc or SARS-CoV-2-RBD mut-hFc protein (5 μg/ml) was added and incubated at 37 °C for 0.5 h. After washing, a goat anti-human Fcγ HRP antibody (Jackson ImmunoResearch, West Grove, PA) was added and incubated at 37 °C for 0.5 h. TMB and H_2_SO_4_ were added to detect the OD_450_ nm value.

### Cell binding and antibody blocking assays

For the cell binding assay, cell suspension of SARS-CoV-2-spike-CHO cells was incubated with the indicated antibody (5 μg/ml) for 1 h on ice and then incubated with goat anti-human PE antibody (Thermo, Pudong New Area, Shanghai) for 1 h on ice. The cells were analyzed by FACS Calibur (BD Biosciences, San Jose, CA). For the antibody blocking assay, antibodies were preincubated with 2.5 μg/ml SARS-CoV-1-RBD-hFc or SARS-CoV-2-RBD-hFc at different concentrations for 1 h on ice, and then, the mixture was incubated with ACE2-CHO cells for 1 h on ice. After washing, the cell suspension was labeled with goat anti-human PE antibody (Thermo, Pudong New Area, Shanghai) and incubated for 1 h on ice. The cells were analyzed by FACS Calibur (BD Biosciences, San Jose, CA).

### Surface plasmon resonance

Surface plasmon resonance (SPR) was performed by GenScript (GenScript, Nanjing) to measure the affinity of the antibody. Antibodies were immobilized on the Series S Sensor Chip Protein A chip (GE Healthcare), and then SARS-CoV-2-RBD-his protein with a gradient concentration from 1.25 nM to 40 nM was injected into the chip. The analysis was performed at a constant temperature of 25 °C. The buffer was HBS-EP +: 10 mM HEPES, 150 mM NaCl, 3 mM EDTA, 0.05% P20, pH 7.4 (lot No. 30393) (GE Healthcare); the flow rate was 10 μl/min. The assay was performed by Biacore T200, GR18010468 (GE Healthcare). Calculation of the combined kinetic constants was performed by Biacore T200 Evaluation software version 3.1.

### Pseudovirus neutralization assay

To generate SARS-CoV-2 pseudovirus, we replaced the coding sequence of VSV-G protein with the sequence of SARS-CoV-2 spike in lentiviral packaging system^19,20^ and then co-transfect HEK293T cells with the pLVX-EGFP-Luciferase reporter gene. The pseudovirus supernatant was collected 48 h later and titrated to 10^5^ pfu/ml. Neutralization assays were performed by incubating pseudovirus with a series of diluted antibodies at 37 °C for 1 h. Then, the pseudovirus-antibody mixture was added to seeded ACE2-CHO cells (approximately 5×10^3^ pfu vs 10^4^ cells/well) in 96-well plates. The half-maximal inhibitory concentration (IC_50_) of each antibody was determined by measuring luciferase activity 48 h later.

### Live SARS-CoV-2 neutralization assay

All experiments about live SARS-CoV-2 were performed under the approved standard operating procedures of Biosafety Level 3 laboratory. Live SARS-CoV-2 was isolated from throat swabs of SARS-CoV-2-infected patients in Jiangsu Province and identified by sequencing (strai Beta CoV-JS27). Viruses were amplified in Vero E6 cells and made as working stocks at 10^5^ pfu/ml. For the neutralization assay, Vero E6 cells were seeded into 96-well plates at 10^4^/well and cultured overnight. SARS-CoV-2 (100 TCID50) was pre-incubated with a series of diluted antibodies at 37 °C for 1 h. Then, the virus-antibody mixtures were added to seeded Vero E6 cells. Cytopathic effects (CPEs) were photographed 4 days later.

### Statistical analysis

All group data are expressed as the mean ± standard deviation (SD) of a representative experiment performed at least in triplicate, and similar results were obtained in at least three independent experiments. All statistical analyses were conducted using GraphPad Prism 8.0. Two-tailed Student’s t-test of the means was used for statistical analysis, with *P* *<0.05 defined as significant.

## Results

### Purification of SARS-CoV-2-RBD mut with a disrupted ACE2 interface as the negative antigen

To screen potent neutralizing antibodies against SARS-CoV-2, we first analyzed the ACE2 interface of SARS-CoV-2-RBD (Figure 1A). We selected sixteen residues essential for the hydrophobic or electrostatic effects within the ACE2 interface of SARS-CoV-2 and made mutations (Table 1). The predicted structure model of SARS-CoV-2-RBD mut showed that the overall conformation of the RBD did not change after mutation (Figure 1B), but the surface property of the ACE2 interface had been changed (Figure 1C). We then fused SARS-CoV-2-RBD, SARS-CoV-2-RBD mut and SARS-CoV-1-RBD with a human Fc tag and performed purification (Figure 1D). The ACE2-binding activity of purified SARS-CoV-2-RBD mut significantly decreased compared to that of SARS-CoV-2-RBD (Figure 1E), and a similar trend was also observed when we detected the binding of SARS-CoV-2-RBD and SARS-CoV-2-RBD mut on ACE2-CHO cells (Figure 1F and Figure 1G). These results indicated that these mutations successfully abolished ACE2 recognition by destroyed ACE2 interface of SARS-CoV-2-RBD. The purified SARS-CoV-2-RBD mut would be suitable to function as the negative antigen in our screening.

**Figure 1.**
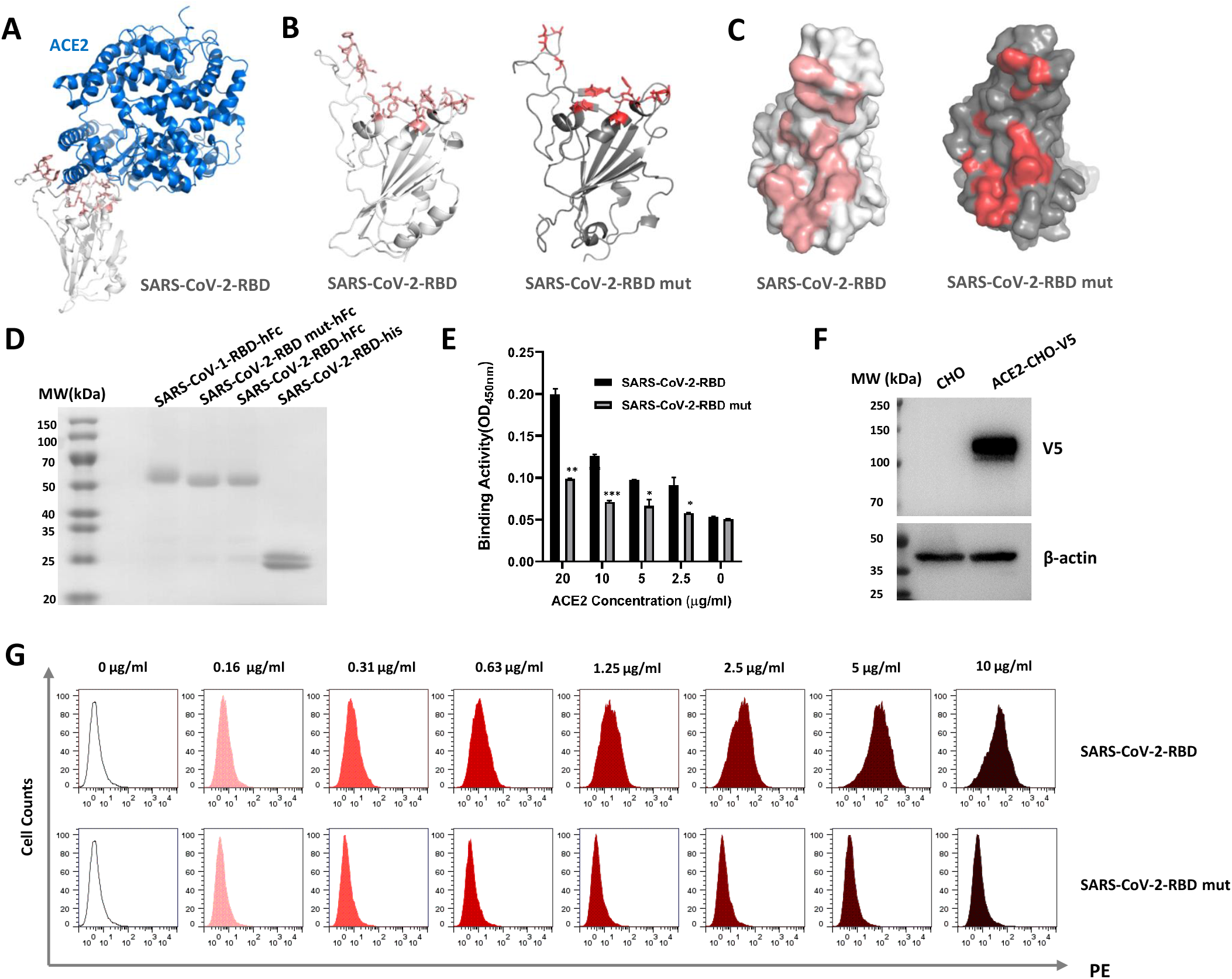
Mutating the ACE2 interface of SARS-CoV-2-RBD to abolish ACE2 binding. (A). Structural model of the SARS-CoV-2-RBD/ACE2 complex. ACE2 interface of SARS-CoV-2-RBD: light red. (B). ACE2-interface comparison for SARS-CoV-2-RBD (light red stick) and SARS-CoV-2 RBD mut (dark red stick). (C). Surface exhibition of the ACE2 interface. (D). SDS-PAGE of purified RBDs. (E). ELISA to detect the ACE2-binding activity of SARS-CoV-2-RBD and SARS-CoV-2-RBD mut. Values represent the mean ± SD, with *P* *<0.05, *P* **<0.01 and *P* ***<0.001 (F). Western blotting to detect ACE2 expression in ACE2-overexpressing CHO cells. (G) Flow cytometry to detect the binding activity of SARS-CoV-2-RBD and SARS-CoV-2-RBD mut on ACE2-CHO cells.

### Isolating SARS-CoV-2-specific antibodies by *in vitro* site-directed screening

To obtain SARS-CoV-2-specific neutralizing antibody, we performed site-directed antibody screening by phage display. We utilized SARS-CoV-2-RBD-his and SARS-CoV-2-RBD-hFc as the positive antigens and GPC5-his and SARS-CoV-2-RBD mut-hFc as the negative antigens to execute the selection within a naive human scFv antibody phage library and a domain antibody phage library, respectively (Figure 2A). After four rounds of screening, the antigen-binding activity of the eluted phage dramatically increased (Figure 2B). Notably, the eluted phage exhibited a stronger binding signal on SARS-CoV-2-RBD compared to that on SARS-CoV-2-RBD mut, especially those from the domain antibody library (Figure 2C), indicating an expected precleaning effect during selection. We then randomly picked 200 single clones from the 4^th^ round eluted phage and performed monoclonal phage ELISA. The positive binders were enriched significantly in both libraries (Figure 2D). All the positive binders were sequenced, and finally, we obtained nine enriched clones from the domain antibody library and one enriched clone from the scFv antibody library. Among them, nine domain antibodies bound to SARS-CoV-2-RBD specifically, whereas one scFv antibody (4A3) showed weak binding activity on SARS-CoV-1-RBD and SARS-CoV-2-RBD mut in the phage ELISA (Figure 2E).

**Figure 2.**
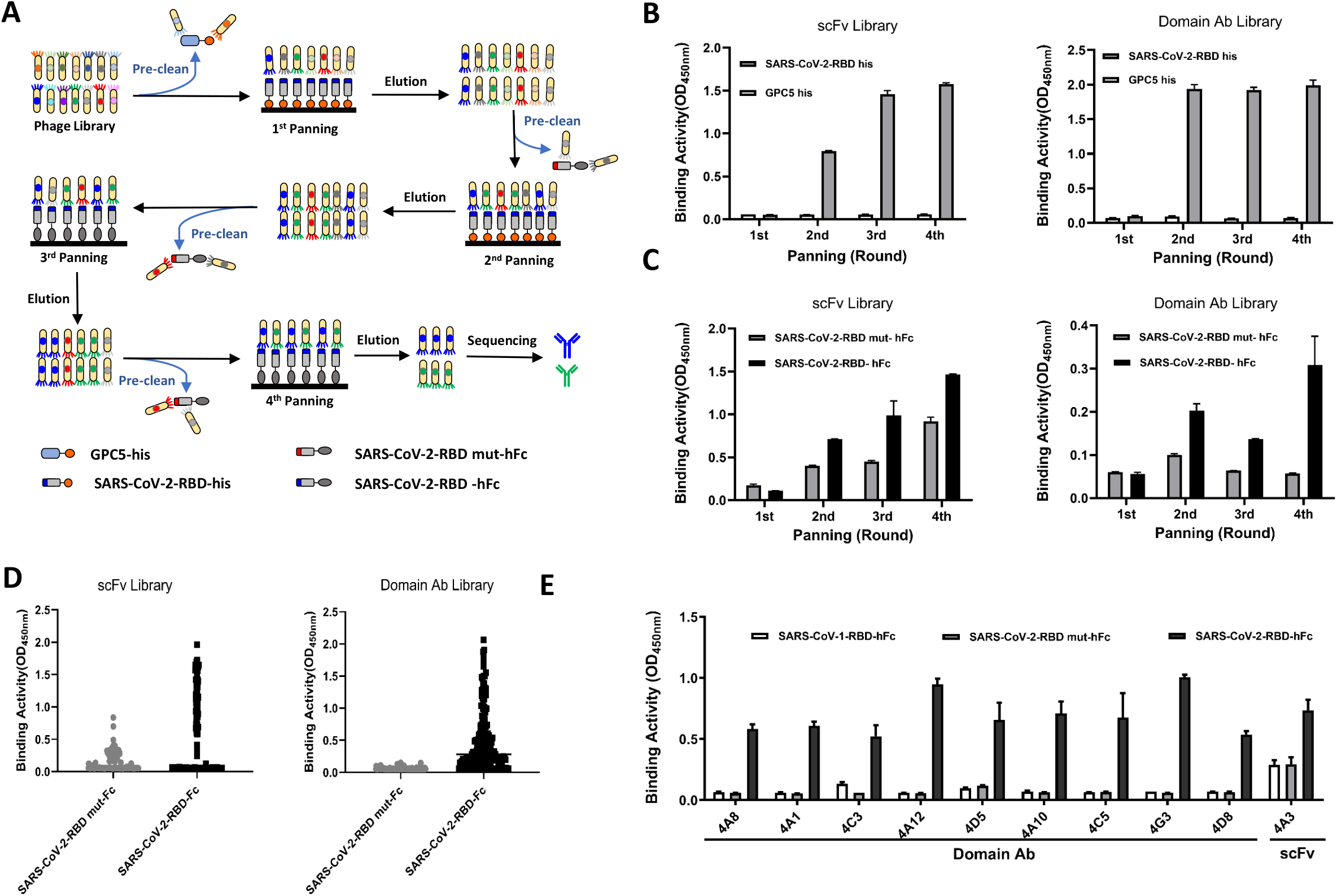
Antibody screening by site-directed phage display. A. Schematic diagram of the screening. Positive antigens: SARS-CoV-2-RBD-his and SARS-CoV-2-RBD-hFc; negative antigens: GPC5-his and SARS-CoV-2-RBD mut-hFc. B. Polyclonal phage ELISA to detect the antigen-binding activity of four rounds of rescued phages. GPC5-his was used as a negative antigen control. C. Polyclonal phage ELISA to compare the binding activities of the eluted phage for SARS-CoV-2-RBD-hFc and SARS-CoV-2-RBD mut-hFc. D. Monoclonal phage ELISA to analyze SARS-CoV-2-RBD-specific binders. E. Capture phage ELISA to detect the antigen-binding specificity of soluble antibodies extracted from TG1 periplasm.

The nine binders isolated from the domain antibody library contained only the antibody heavy chain variable region. We then fused them with a human Fc tag and performed purification. The 4A3 scFv binder was converted into a human IgG1 and purified as well (Figure 3A). Among all the purified antibodies, 4A12, 4D5, 4A10, 4C5 and 4A3 were selected for further evaluation due to their strong binding activities to both SARS-CoV-2-RBD (Figure 3B) and SARS-CoV-2 spike-overexpressing cells (Figure 3C), in addition to their promising expression yield and purity.

**Figure 3.**
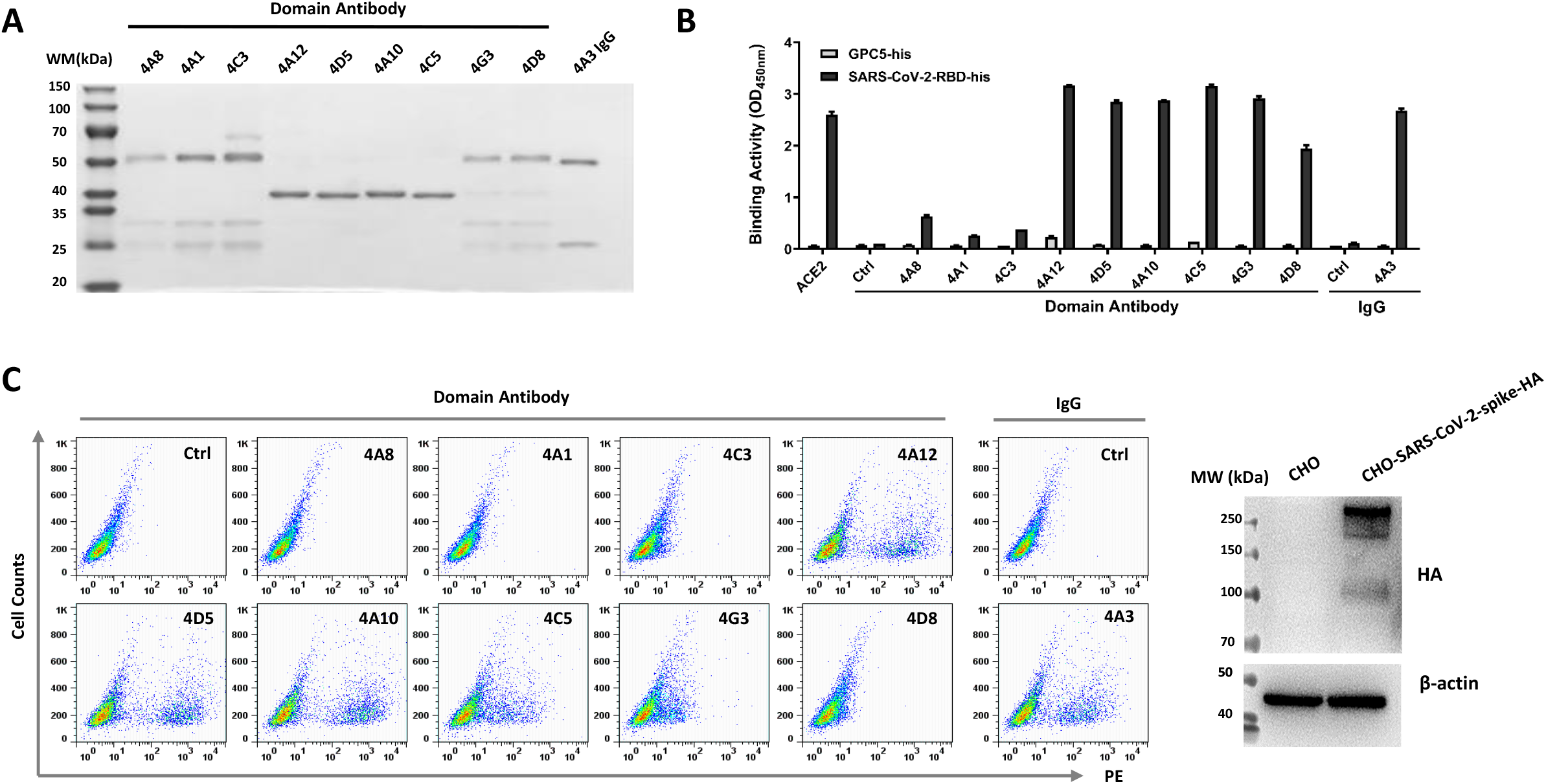
Binding properties of purified antibodies on SARS-CoV-2-RBD and SARS-CoV-2-spike-overexpressing cells. (A). SDS-PAGE of purified antibodies. (B). ELISA to detect the binding activity of the purified antibodies for SARS-CoV-2-RBD-his. GPC5-his was used as a negative antigen control. (C). Flow cytometry to detect the binding activity of the purified antibodies on SARS-CoV-2-spike-CHO cells (left). Western blot showing the expression level of spike in SARS-CoV-2-spike-CHO cells (right).

### The candidate antibodies blocked the binding of SARS-CoV-2-RBD to ACE2positive cells

To examine the potential neutralizing capabilities of our candidate antibodies, we detected whether they would disturb the binding between SARS-CoV-2-RBD and ACE2-positive cells. Three domain antibodies (4A12, 4D5, 4A10) and 4A3 IgG exhibited obvious inhibition in a dose-dependent manner, whereas the SARS-CoV-1-neutralizing antibody M396 seemed to have no effect (Figure 4A and 4C). Considering the high similarity of the RBDs, we also evaluated the blocking effects of our antibodies on the binding between SARS-CoV-1-RBD and ACE2-CHO cells. None of our candidate antibodies showed inhibitory effects (Figure 4B and 4D). Notably, we suspected that clone 4A3, which showed weak cross-reaction with SARS-CoV-1-RBD in the phage ELISA (Figure 2E), might exhibit a certain blocking effect for the cell binding of SARS-CoV-1-RBD, but actually, it showed specific blocking on only the cell binding of SARS-CoV-2-RBD. We then performed SPR to further evaluate the affinities of these candidate antibodies. The results showed that the affinities of our antibodies ranged from 1.03 nM to 5.82 nM (Table 2 and Figure 5). Altogether, these results indicated that we obtained four antibodies that might have potential neutralizing functions against SARS-CoV-2.

**Figure 4.**
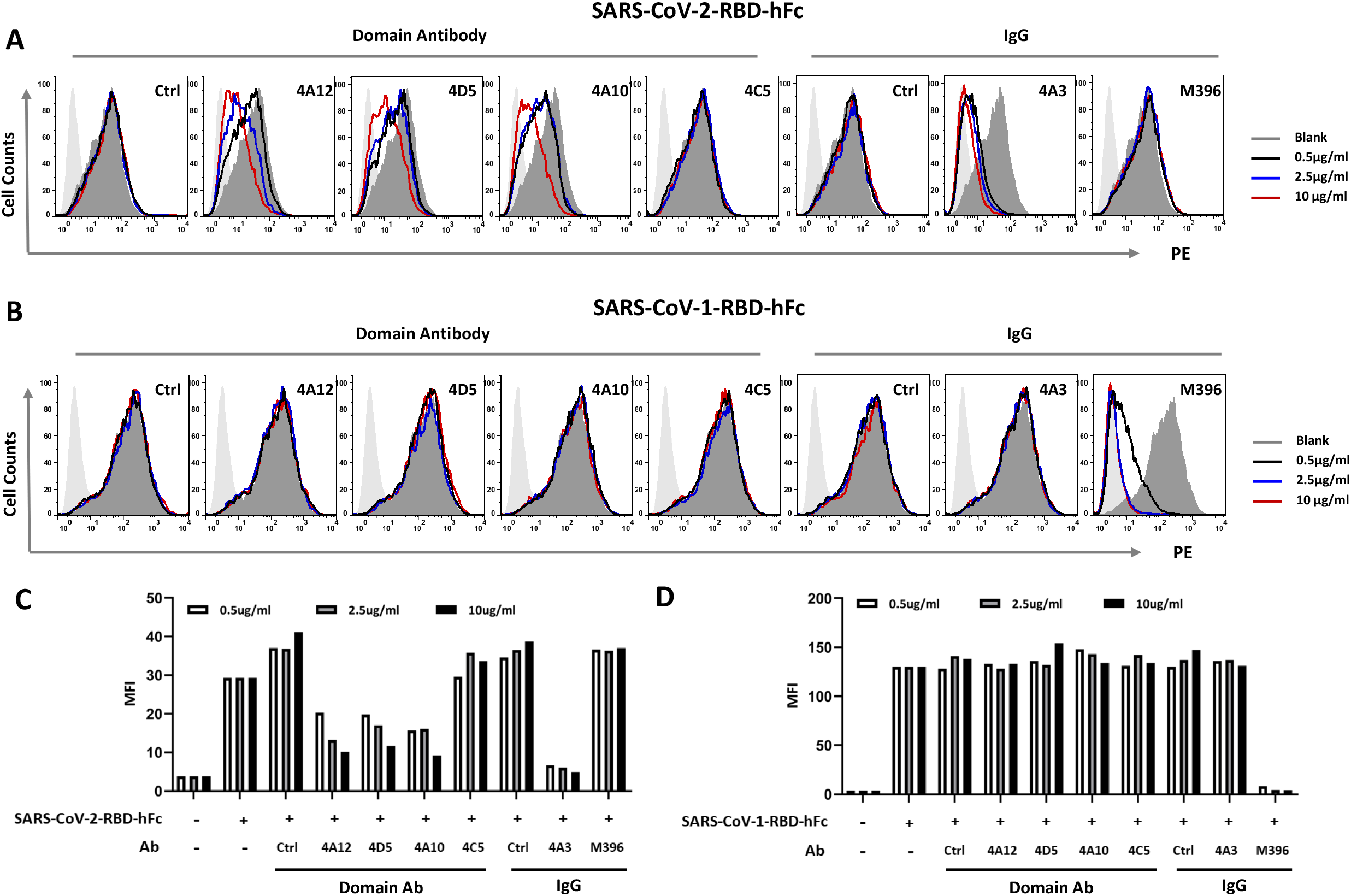
Several selected antibodies blocked SARS-CoV-2-RBD binding to ACE2-CHO cells. (A). Flow cytometry to examine the blocking effect of selected antibodies on the binding of SARS-CoV-2-RBD-hFc and CHO-ACE2 cells. M396, the neutralizing antibody against SARS-CoV-1 was used as a negative antibody control. The mean fluorescence intensity (MFI) of the binding is shown as a histogram in (C). (B). Flow cytometry to examine the blocking effect of selected antibodies on the binding of SARS-CoV-1-RBD-hFc and CHO-ACE2 cells. M396 was used as a positive antibody control. The MFI of the binding is shown as a histogram in (D).

**Figure 5.**
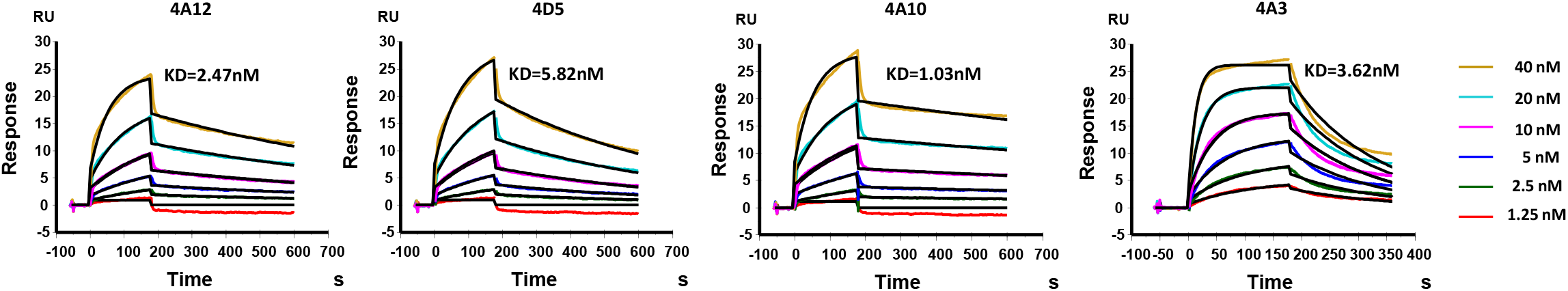
Antibody affinity measurement and spike trimer-binding prediction. SPR to measure the binding affinity of SARS-CoV-2-RBD-his protein to captured antibodies.

**Table 2.** Affinity measurement of RBD protein to antibodies.

| Ligand | Analyte | *ka* (1/Ms) | *kd* (1/s) | KD (M) | Rmax (RU) | Chi<sup>2</sup> (RU<sup>2</sup>) |
| --- | --- | --- | --- | --- | --- | --- |
| 4A3 IgG | RBD protein | 1.91E+06 | 6.90E-03 | 3.62E-09 | 25.55 | 0.379 |
| 4A10 | RBD protein | 4.48E+05 | 4.61E-04 | 1.03E-09 | 20.86 | 0.38 |
| 4A12 | RBD protein | 4.29E+05 | 1.06E-03 | 2.47E-09 | 18.52 | 0.302 |
| 4D5 | RBD protein | 2.98E+05 | 1.74E-03 | 5.82E-09 | 24.33 | 0.304 |

### Domain antibody 4A12 was predicted to have advantage for accessing all three ACE2 interfaces of the spike homotrimer

According to the recently reported cryo-EM structure, the SARS-CoV-2 spike trimer appears in two distinct confirmation states: a closed state with the three RBDs embedded and an open state with only one RBD extended for ACE2 binding. Since the extended RBD presents a complete exposed ACE2 interface that would be more easily captured by an antibody than the closed RBD, we simply wondered whether the remaining embedded ACE2 interfaces of both the open and closed spike trimers could be accessed by antibody. Because the reported cryo-EM structure of the SARS-CoV-2 spike trimer lacks some RBD residue information, we replaced its RBD with the determined structure of SARS-CoV-2-RBD to establish the remodeled SARS-CoV-2 spike trimer. We then analyzed the interspace between each embedded ACE2 interface and its neighboring monomer. The three interspaces in the closed trimer were quite uniform (from 17.2 Å to 19.5 Å). For the two interspaces of the open trimer, one did not change (19.5 Å), whereas the other was somehow occupied by the extended RBD (11.2 Å) (Figure 6A). The domain antibody exhibited a relatively smaller size and antigen interface because it contains only a heavy chain variable region with three complementarity-determining regions (CDRs) instead of six in IgG (Figure 6B). We then performed molecular docking to compare the binding patterns of 4A3 scFv and the domain antibodies on our remodeled spike trimers. Both 4A3 and the domain antibodies were predicted to recognize all three ACE2 interfaces of the closed trimer (Figure 6C). However, when docked with the open trimer, 4A3 scFv was predicted to target only two ACE2 interfaces. It could not access the embedded ACE2 interface occupied by the extended RBD, probably due to its larger size. Surprisingly, one of the domain antibodies, 4A12, was predicted to block all three ACE2 interfaces (Figure 6D). These predictions suggested that the domain antibody might have the advantage of blocking all the ACE2 interfaces of both closed and open trimers, which might offer more effective protection against SARS-CoV-2. Nevertheless, detailed structural examination of the binding patterns of these antibodies with SARS-CoV-2-RBD and SARS-CoV-2 spike trimers is extremely important in our future investigations.

**Figure 6.**
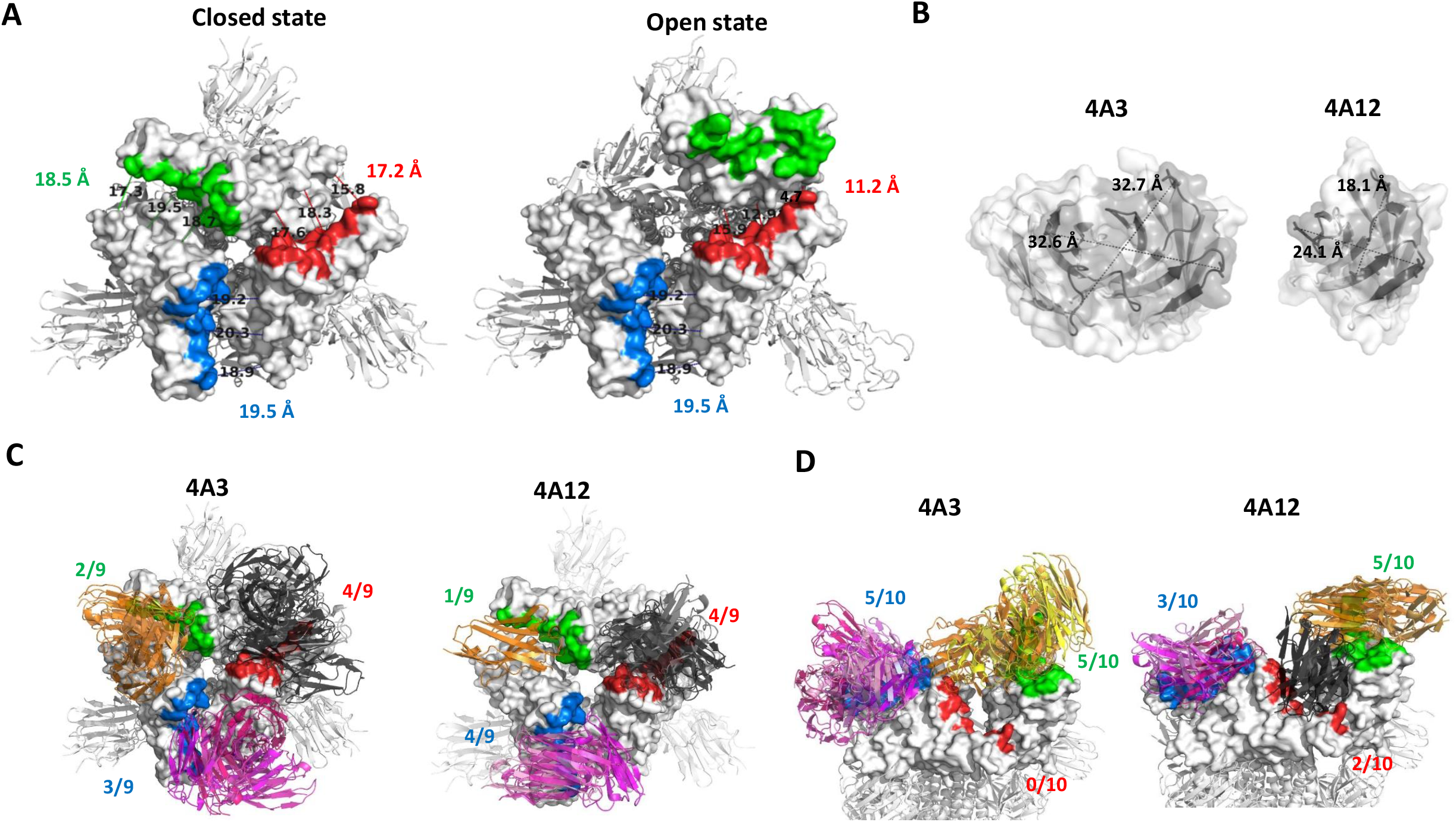
The binding patterns of 4A3 and the domain antibody 4A12 on modeled SARS-CoV-2 spike trimers. (A). The modeled structures of SARS-CoV-2 spike trimers. The RBDs are shown on the surface, and the ACE2 interface of each monomer is labeled in color. The interspace between each ACE2 interface and its neighboring monomer is shown as the average distance of three measurements. (B). The CDR areas of 4A3 scFv and the domain antibody 4A12 (dark). The largest section of each CDR is shown as the indicated length and width. (C). The docking complexes of 4A3 scFv/closed SARS-CoV-2 spike trimer (left) and domain antibody 4A12/closed SARS-CoV-2 spike trimer (right). Antibodies are shown in cartoon. The top nine predictions are shown for each antibody. (D). The docking complexes of 4A3 scFv/open SARS-CoV-2 spike trimer (left) and domain antibody 4A12/open SARS-CoV-2 spike trimer (right). Antibodies are shown in cartoon. The top 10 predictions are shown for each antibody.

### The candidate antibodies exhibited potent neutralizing activities against SARS-CoV-2 pseudovirus and authentic SARS-CoV-2 virus

To evaluate the potential antiviral activities of our antibodies, we prepared SARS-CoV-2 pseudovirus by replacing the coding sequence of VSV glycoprotein with SARS-CoV-2 spike glycoprotein in a lentivirus packaging system (Figure 7A). We preincubated our candidate antibodies with SARS-CoV-2 pseudovirus and then added them to cultured ACE2-CHO cells. The domain antibodies 4A12, 4A10, and 4D5 and 4A3 IgG exhibited obvious neutralizing potencies, with IC_50_ values from 0.19 μg/ml to 1.13 μg/ml (Figure 7B). These results were consistent with the blocking pattern observed in the SARS-CoV-2-RBD and ACE2-CHO cell-binding assays (Figure 4A and 4C). Although 4C5 did not show a blocking effect in our cell-binding assay, it still showed a mild neutralizing capability (Figure 7B).

**Figure 7.**
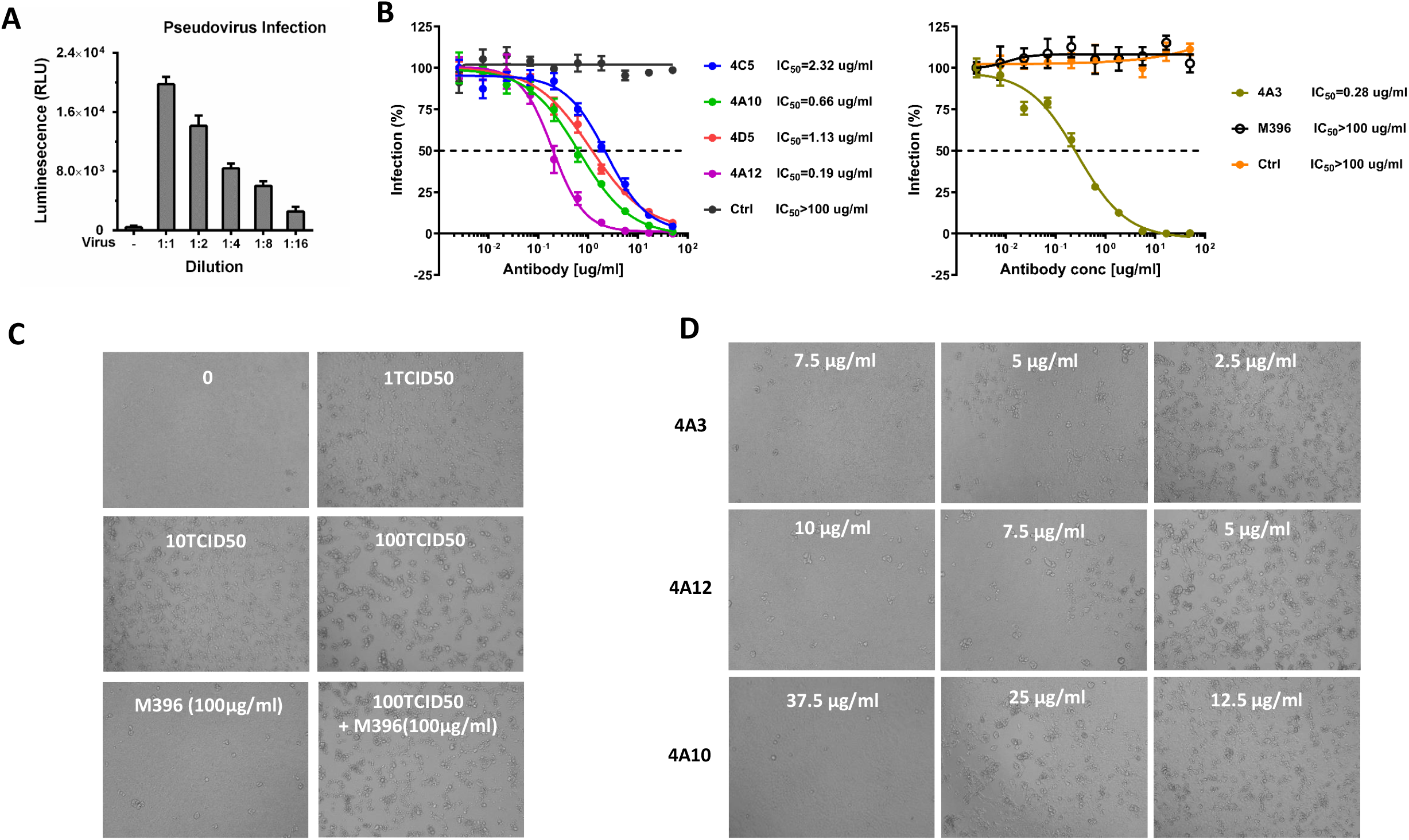
Antibody neutralization analyzed by pseudovirus and live SARS-CoV-2. (A) Quality control of SARS-CoV-2 pseudovirus. Luciferase reporter assay to detect SARS-CoV-2 pseudovirus infection in CHO-ACE2 cells. (B) Neutralization effects of candidate antibodies on SARS-CoV-2 pseudovirus were analyzed by infecting CHO-ACE2 cells with antibody-blocked SARS-CoV-2 pseudovirus. The domain antibody 31A2 against galectin-3 was used as the control domain antibody. 32A9 IgG targeting glypican-3 and the SARS-CoV-1-RBD-specific neutralizing antibody M396 were used as control antibodies for 4A3 IgG. (C). Quality control of live SARS-CoV-2. Photographed CPE of Vero E6 cells exposed to 1 TCID50, 10 TCID50 and 100 TCID50 of SARS-CoV-2 for 4 days. (D). Photographed CPE to show the neutralization effects of candidate antibodies by infecting Vero E6 cells with antibody-blocked SARS-CoV-2 (100 TCID50) for 4 days.

Based on our pseudovirus experiment, we selected 4A3 IgG, 4A12 and 4A10 to evaluate their neutralizing activities on authentic SARS-CoV-2. The observations of CPE showed that 4A3, 4A12 and 4A10 exhibited complete protection at 7.5 μg/ml, 10 μg/ml and 37.5 μg/ml, respectively, when we performed a 4-day exposure to SARS-CoV-2 (100 TCID50) (Figure 7C and 7D). These protective effects were still quite stable when the exposure time was extended to 10 to 15 days (Table 3).

**Table 3.**
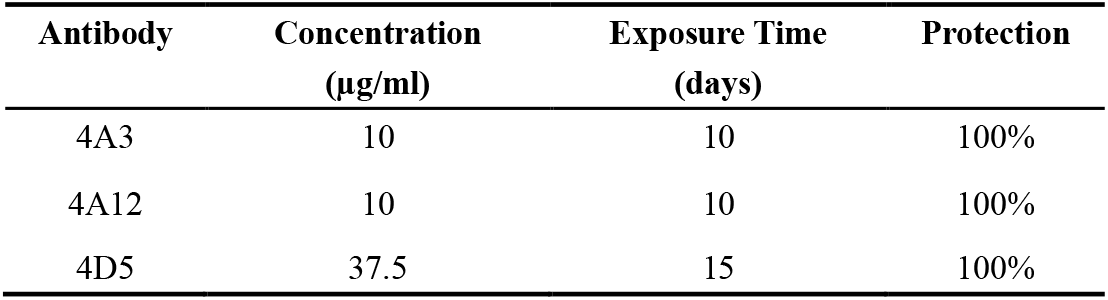
Long-term SARS-CoV-2-neutralizing effects of candidate antibodies.

| **Antibody** | **Concentration**<br>**($\mu\text{g/ml}$)** | **Exposure Time**<br>**(days)** | **Protection** |
| --- | --- | --- | --- |
| 4A3 | 10 | 10 | 100% |
| 4A12 | 10 | 10 | 100% |
| 4D5 | 37.5 | 15 | 100% |

Overall, we isolated several human monoclonal neutralizing antibodies against SARS-CoV-2 by site-directed screening strategy, which could be promising candidate drugs for the prevention and treatment of COVID-19.

## Discussion

Currently, several studies have obtained SARS-CoV-2 antibodies with neutralizing activity by phage display, including full length antibody^21^ and domain antibody^22,23^. Instead of using a mutant SARS-CoV-2-RBD with a disrupted ACE2 binding motif, one of these studies utilizing a captured ACE2/SARS-CoV-2-RBD complex to perform the pre-absorption which also obtained neutralizing antibodies as well^21^.

The RBDs mediate the ACE2 interaction for both SARS-CoV-1 and SARS-CoV-2^13^. Many investigations utilize the RBD as the target region to produce neutralizing antibodies and vaccines^24–26^. It has been reported that immunizing rodents with SARS-CoV-2-RBD elicits a robust neutralizing antibody response without antibody-dependent enhancement (ADE)^27^. According to the determined structure, the ACE2 interface of the RBD presents only a small portion of the whole domain^15^, and only those antibodies binding to the interface would directly interfere the interaction with ACE2. This would be one possible explanation for why not all the antibodies isolated with RBD binding activity showed neutralizing activity. Based on this information, we performed site-directed screening with the positive antigen SARS-CoV-2-RBD and negative antigen SARS-CoV-2-RBD mut to ensure that we obtained antibodies accurately targeting the ACE2 interface. As we predicted, most of our candidate antibodies exhibited a significant blocking effect for ACE2 recognition on cells (Figure 4). Although the antibody 4C5 did not, it did show a relatively mild virus-neutralizing function compared to that of the other candidates (Figure 7). These functional evaluations might prove the feasibility of our site-directed screening strategy, but further validations, such as epitope determination by cryo-EM, are still necessary.

The small size of the domain antibody enables several unique advantages, including high expression yield, enhanced tissue penetration, and hidden epitope targeting^28–30^. The structural transition from the closed to open state of the RBD has been proven necessary for receptor engagement and membrane fusion in coronaviruses^31,32^. Therefore, blocking all the RBDs, especially all the ACE2 interfaces of both open and closed spike trimers, by antibodies would theoretically offer enhanced protection against virus infection. It seemed that one of our domain antibodies, 4A12, might execute its neutralizing function in this way, but additional structural evidence is needed. These observations also suggest that the neutralizing mechanism of antibody should be evaluated on both SARS-CoV-2-RBD and SARS-CoV-2 spike trimer. Because a recent study reports that the neutralizing activity of CR3022, a SARS-CoV-1 neutralizing antibody, is lost on SARS-CoV-2 due to the different conformation characters of spike trimer^33^. On the other hand, a smaller size causes a shorter half-life of the domain antibody *in vivo*^34,35^, which might be a potential disadvantage for its neutralizing function. However, one of the major safety concerns of coronavirus vaccines is the undesirable ADE induced by anti-spike antibodies^36,37^. Although several studies have demonstrated that neutralizing antibodies seem less likely to induce ADE than nonneutralizing antibodies^38–40^, we could not exclude possible ADE since there is still no available clinical result for either a SARS-CoV-2 vaccine or neutralizing antibody. In this case, a domain antibody with a shorter half-life would be a possible choice to balance neutralization and ADE side effects.

Phage display is a well-established strategy for *in vitro* antibody screening^41^. The antibody is amplified in the prokaryotic system quickly and efficiently^42^. We initially obtained ten enriched binders that all showed strong antigen-binding activity. However, when expressed in 293T cells, three of them lost antigen binding, and two of them exhibited poor yield. This effect is a common problem for phage display when shifting to a eukaryotic expression system. Other *in vitro* screening strategies, such as yeast display^43^ or mammalian cell display^44^, may offer more stable selection due to their eukaryotic background. Antibodies selected from naive phage libraries usually require further optimization to improve their affinity. Even though the affinities of our antibodies are in a range (10^-9^ M) similar to that of antibodies screened from SARS-CoV-2-infected patients^11^, we still need to perform affinity maturation to achieve more competitive affinity and neutralizing function against SARS-CoV-2. On the other hand, collecting the blood of infected donors to construct a SARS-CoV-2-specific phage library for antibody screening will be an alternative strategy as well.

In the current study, we established site-directed phage display screening, a feasible and efficient *in vitro* assay, to obtain neutralizing antibodies. In addition to screen neutralizing antibodies against viruses, this strategy could be widely used for isolating function-blocking antibodies targeting the essential domains of various antigens. Moreover, the SARS-CoV-2-neutralizing antibodies isolated here presented in both IgG and single domain forms, which could offer valuable research tools for understanding the mechanisms of SARS-CoV-2 pathogenesis, such as virus-host recognition and cell entry. Most importantly, these antibodies with validated neutralizing function against SARS-CoV-2 could be promising drugs for the clinical application of COVID-19 prevention and treatment.

## Acknowledgments

We thank our colleagues Dr. Yujie Sun and Dr. Ningning Wang for providing cell lines; Dr. Fan Lin and Dr. Shuo Yang for helping with the pseudovirus detection assays; and Dr. Yujie Sun, Dr. Tong Ding and Dr. Luan Sun for suggestions. This research was supported by the National Natural Science Foundation of China (No. 81773260 and No. 81972284), National Science and Technology Major Project (2018ZX10734401-006), and National Natural Science Foundation Youth Project of Jiangsu, China (No. BK20171047).

## Authors’ Contributions

XL, FG, LG, YC, LA and YG performed the experiments; XL, FG, LG, XG and WG analyzed the data; WG, XG, ZH and HS designed and conducted the research; WG and XL wrote the manuscript; and XG, ZH and HS revised the manuscript. All authors read and approved the final manuscript.

## Competing interests

The authors declare that they have no conflicts of interest with the contents of this article.

## Notes

### Competing Interest Statement

The authors have declared no competing interest.

### Summary of Updates

The title has been changed.

